# Trips and Neurotransmitters: Discovering Principled Patterns across 6,850 Hallucinogenic Experiences

**DOI:** 10.1101/2021.07.13.452263

**Authors:** Galen Ballentine, Samuel Freesun Friedman, Danilo Bzdok

**Author notes:** equal contribution.

## Abstract

Psychedelics are thought to alter states of consciousness by disrupting how the higher association cortex governs bottom-up sensory signals. Individual hallucinogenic drugs are usually studied in participants in controlled laboratory settings. Here, we have explored word usage in 6,850 free-form testimonials with 27 drugs through the prism of 40 neurotransmitter receptor subtypes, which were then mapped to 3D coordinates in the brain via their gene transcription levels from invasive tissue probes. Despite the variable subjective nature of hallucinogenic experiences, our pattern-learning approach delineated how drug-induced changes of conscious awareness (e.g., dissolving self-world boundaries or fractal distortion of visual perception) are linked to cortex-wide anatomical distributions of receptor density proxies. The dominant explanatory factor related ego-dissolution-like phenomena to a constellation of 5-HT2A, D2, KOR, and NMDA receptors, anchored especially in the brain’s deep hierarchy (epitomized by the associative higher-order cortex) and shallow hierarchy (epitomized by the visual cortex). Additional factors captured psychological phenomena in which emotions (5-HT2A and Imidazoline1) were in tension with auditory (SERT, 5-HT1A) or visual (5-HT2A) sensations. Each discovered receptor-experience factor spanned between a higher-level association pole and a sensory input pole, which may relate to the previously reported collapse of hierarchical order among large-scale networks. Simultaneously considering many psychoactive molecules and thousands of natural language descriptions of drug experiences our framework finds the underlying semantic structure and maps it directly to the brain. These advances could assist in unlocking their wide-ranging potential for medical treatment.

## INTRODUCTION

For thousands of years, humans have been drawn to consume hallucinogenic substances to deliberately alter states of consciousness. These drug-induced mental states frequently involve mystical experiences, the dissolution of boundaries between self and world, changes of social-emotional perception, and a dramatic intensification of sensoria^1^. Mind expansion experiences have served as vehicles for religious ceremonies, oracle traditions, and healing rituals^2–4^. The subjective alterations of reality are known to be highly variable across individuals. This variability of the nature of drug-induced experiences may depend on one’s life history, world view, and the setting of the experience^5^.

Inter-individual variability poses a key challenge as we venture to bring hallucinogenic substances into medical practice. The same drug can induce boundless feelings of joy and love in some sessions, but terror and panic in other sessions^6–8^. In a rapidly growing number of successful clinical studies, psychedelic drugs show promise as treatments for a variety of psychiatric conditions^9–12^. As one recent example, a clinical trial^13^ examined psilocybin-assisted psychotherapy for treatment-resistant major depression (the active compound of “magic mushrooms”). After only two drug doses, investigators found a large and durable treatment benefit in 71% of participants at 8 weeks after the intervention.

Such clinical studies have been accompanied by some early investigations into the neurophysiological basis that underlies the drug effects on high-level thought^14–16^. Crucially, the efficacy of these substances to treat mental illness has been tightly linked to specific experiential phenomena such as ‘ego dissolution’^17–19^. Hence, it is now urgent to systematize the nuances of the caused changes in states of conscious awareness and the molecular mechanisms that relate to neural processes that underpin them.

The ‘classical psychedelics’ are thought to alter perception of self and the world by acting primarily on the 5-HT2A receptor. This neurotransmitter receptor is targeted by LSD, psilocybin, mescaline, DMT, and phenethylamines. For several decades, pharmacological research has placed a strong focus on 5-HT2A as a putative essential mechanism of hallucinogenic experiences^20^. In contrast to the idea of selective stimulation of the 5-HT2A receptor, debate has recently emerged as to whether the differences in experience induced by different drugs are explained by functional selectivity at the 5HT2a receptor itself or, in contrast, orchestrated by the vast array of neurotransmitter receptor subclasses on which these drugs act^1,21^.

In support of this latter notion, similar alterations in subjective awareness have been reported by hallucinogenic drugs that are biochemically and pharmacologically distinct from the classical psychedelics. Examples of these non-traditional psychedelic-like drugs include ketamine (i.e., NMDA-antagonist and dissociative anaesthetic), MDMA (an “entactogenic” amphetamine-derivative), ibogaine, and salvia divinorum (a novel kappa-opioid agonist). As a key motivation for our present investigation, such drugs interact minimally, indirectly, or not at all with 5-HT2A receptors in the brain^22–24^. This fact hints at a richer spectrum of neurobiological mechanisms underlying “psychedelic” action.

We will henceforth use the umbrella term ‘hallucinogen’ to refer to compounds that have been reported to temporarily modify conscious awareness. There is overlap and idiosyncrasy in the target receptor profiles of hallucinogens. Additionally, such compounds cause both unique and convergent effects on subjective perception. This circumstance calls for new research avenues that can illuminate the common denominator across a variety of hallucinogenic experiences and ingested drugs. Such an approach has the potential to shed light on general principles by which receptor binding patterns trigger specific forms of subjective experience – a goal that is hard to reach with the sample sizes typical in today’s clinical trials on hallucinogens.

On the macroscopic level, hallucinogens are thought to mediate psychological effects through functional coupling shifts between large-scale brain networks. Especially, a key role has been ascribed to induced rebalancing between lower sensory processing and higher-level integration^15,25,26^. In particular, hallucinogens have been thought to “reduce the precision of high-level priors (expectations or beliefs about the world) and concurrently increase bottom-up information flow”^27^. Integrative processing across major brain networks probably supports the stable sense of self which experiences sensory stimuli. Triggered by hallucinogens, the resulting neural dynamics may reflect a flood of minimally filtered processing of interoceptive and exteroceptive sensory information, and a concomitant disintegration of ordinary self-awareness^28,29^. This shift in functional interplay between large-scale networks is modulated in the hallucinogenic state through the key receptors that are expressed throughout the cerebral cortex.

A complete understanding of the mechanism of hallucinogenic drug action should ultimately account for both functional activity at receptors as well as the neuroanatomical distribution of these receptors throughout the cortex. While the functional changes induced by binding at these receptors depends on multiple parameters--degree of inhibition, activation, functional selectivity--that are beyond the scope of the present investigation, we aim instead to explore the more circumscribed problem of the neuroanatomical distribution of these receptor binding patterns. Future studies can build upon this neuroanatomical scaffold to incorporate functional parameters at each receptor.

In this study, we have linked detailed text reports to hallucinogenic drug affinity for neurotransmitter receptor subclasses. Our study supplements previous psychedelic research in at least three ways: a) We have extracted knowledge by mining 6,850 real-world experience samples, in which hallucinators freely describe their subjective psychedelic experiences. In contrast, laboratory studies or randomized clinical trials on hallucinogens have been restricted to testing drug effects in a handful of participants. b) Our data-driven approach allowed rigorous testing for the existence of receptor-experience mappings across a diverse array of hallucinogenic drugs. This is a prerequisite for exploring overarching principles that may encapsulate how specific sets of receptor bindings are linked to specific drug-induced alterations of subjective awareness. c) We have aimed to deconstruct the hallucinogenic perturbations of subjective awareness by cutting across a spectrum of molecular receptor subtypes. In an ecologically faithful approach, we describe an anatomic grounding for the subjective experience of altered states of consciousness.

## RESULTS

We aimed to illuminate key principles that mediate hallucinogenic states of consciousness. For this purpose, we have charted regularities across a large corpus of naturalistic psychedelic experiences. Our pattern-learning strategy has dissected phenomenologically rich anecdotes into a ranking of constituent brain-behavior factors: each characterized by a unique neurotransmitter fingerprint of action and a unique experiential context. The cortex-wide distribution of receptor-experience factors was located to anatomical regions across the deepest and shallowest neural network layers. The genetic factor expressions implicated regions, ranging from the higher association cortices to the unimodal sensory cortices. The dominant factor elucidated mystical experience in general, and the dissolution of self-world boundaries (‘ego dissolution’) in particular. The second and third most explanatory factors evoked auditory, visual, and emotional themes of mental expansion. Each subjective drug experience was modeled as a specific combination of the brain-behavior factors. In this way, we have carefully de-constructed the changes of conscious awareness triggered by hallucinogenic drugs into its component parts.

Although not imposed by our analysis approach, the cortical mappings of factor expression often turned out to be spatially contiguous with smooth transitions of expression strength between neighboring brain regions. Clusters of adjacent brain regions with a particular expression strength also respected well-known anatomical and functional divisions. Moreover, salient brain-behavior effects were often found to be mirrored in homologous brain regions in the left and right hemisphere. These observations are noteworthy because all steps of our modeling pipeline are blind to the topographical position of particular brain atlas regions. The results across components were organized for ease of visualization: since the directionality of the poles within each component is arbitrary in CCA, the pole most strongly associated with receptor transcription in the visual cortex was assigned the positive value and listed first.

As per our chosen pattern-learning algorithm (canonical correlation analysis), the strongest factor explained the largest portion of systematic joint variation between receptor affinity profiles and collective experiential terms (explained variance *rho*=0.85, statistically significant at *p* < 0.05; Fig. 1). This dominant receptor-experience factor highlighted a theme of sensory phenomena including terms like *visuals*, *nausea*, *sleep*, *fatigue*, *mild* and similar mentions. Perhaps describing more sensory and visceral qualities of hallucinogenic experience, this constellation can be categorized into interoception (*nausea*, *fatigue*, *sleep*, *stomach*) and exteroception (*vision*, *auditory*, *movie*). These facets of experience reports were most tightly related to the 5-HT2A receptor. However, other serotonin receptors (5-HT2C, 5-HT1A, 5-HT2B), adrenergic receptors Alpha 2a and Beta 2, as well as the D2 receptor also played a role in this experiential theme. This fingerprint of receptor actions was most strongly associated with the drugs DIPT, DOI, and 5-MeO-TMT. Experiences mediated by modulation of these receptor combinations were located at the lower end of the neural processing hierarchy, especially highlighting extended parts of the visual sensory regions.

**Figure 1:**
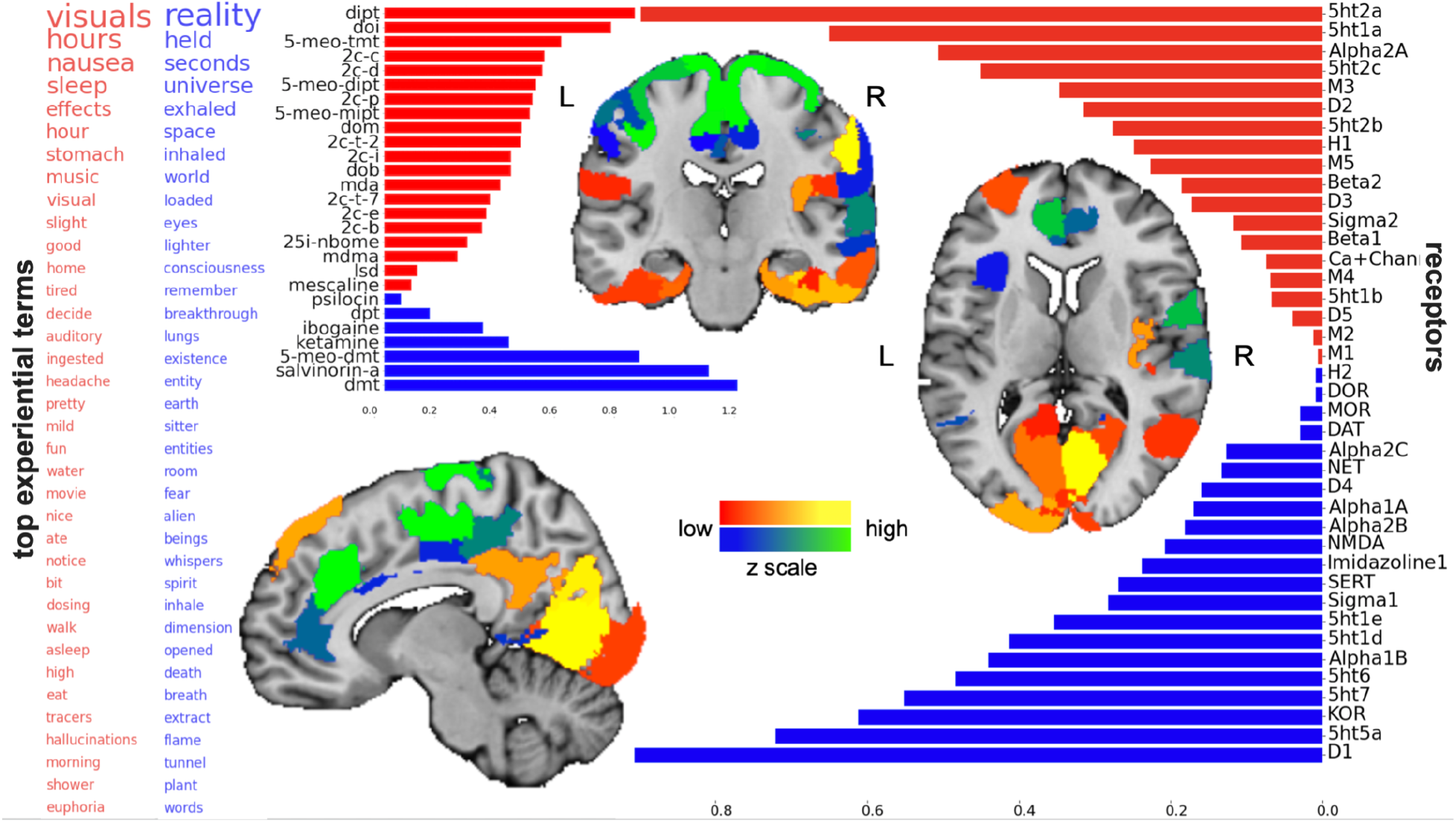
The leading factor underlying hallucinogenic experiences is compatible with the phenomenon of ego dissolution. Shows the statistically strongest pattern of how binding affinities of neurotransmitter receptors were cross-linked with usages of >14,000 words in thousands of real-world hallucinogenic sessions. *Left*: Experiential terms together make apparent the semantic context that was consistent across drugs with diverging pharmacological mechanisms of action. Bigger words had relatively higher ranking in the factor. *Right*: Synaptic receptor density proxies that were most associated with the experiential theme of factor 1. Blue bars correspond to the blue word-values on the left, while red bars correspond to red words. *Middle*: Shows how much each examined drug is associated with the receptor gene profiles of factor 1. Blue bars correspond to the blue word-values on the left, while red bars correspond to red words. Brain renderings highlight those among the 200 anatomical regions where the local co-expression of receptor genes was most tightly linked with factor 1 (NeuroVault link: to_be_added_later). Sagittal, coronal, and axial brain slices are shown at x=+5, y=-14, and z=+15 (MNI reference space). The colors of these renderings demonstrate the density of the weighted receptor gene expression for each region and correspond to the poles of the factor: blue words and bars correspond to blue/green brain regions (green>blue intensity) while red words and bars correspond to red/yellow brain regions (yellow>red intensity).

The dominant factor spanned a continuum of experiential facets whose opposite extreme flagged words that appear to describe mental expansion, including the terms *universe*, *space*, *world*, *consciousness*, *breakthrough*, *existence*, *earth*, *dimension*, *reality*, *flame*, and *tunnel* – all of which would be consistent with the phenomenology of the mystical experience^6^. References to liminal conscious beings, also characteristic of the mystical experience, were described by the terms *entity, sitter, alien, beings*, and *spirit*. A theme of immediate time horizon was indicated by the term *seconds*, and also suggested by references to bodily systems such as *eyes* and *lungs*, along with physiological functions *inhale(d), exhaled*, and *breath*. At the level of neurotransmitter receptors, this constellation of induced conscious alterations was linked most strongly with drugs that preferentially bind to D1, 5-HT7, KOR, 5-HT5A, as well as Sigma 1 and NMDA. The co-dependencies embedded in this profile of receptor bindings and subjective terms were most closely associated with the hallucinogenic drugs DMT, salvinorin A, 5-MeO-DMT, and ketamine. The anatomical brain regions that co-express these sets of receptor density proxies included some of the highest regions of the association cortex, especially the rostral and dorsal anterior cingulate cortex, temporoparietal junction, and also prominently in the primary motor and sensory cortices.

The second most important factor revealed by our pattern-learning algorithm tracked joint variation between delineated receptor affinity profiles and experiential term constellations (explained variance *rho*=0.80, statistically significant at *p* < 0.05; Fig. 2) that were not already detected by factor 1. The separate factor 2 highlighted a theme centered on positive emotions including the terms *friends*, *love*, *wonderful*, *amazing*, *dance*, *dancing*, *magic*, and *happy*, but also negative emotional terms like *depression*, *loss*, and *crying*. These emotionally charged facets of drug-induced experience were associated with report terms that describe an intermediate time range of *days*, *week*, *months*, and *years*. The pattern was linked to a graded affinity for a diverse mix of receptors including especially 5-HT2A and Alpha 2a, which had comparable magnitudes of affinity correlation. Additional but less prominently implicated neurotransmitter receptors included calcium channel, 5-HT2C, M3, 5-HT1B, D2, D5, and 5-HT5A. The joint variation between these receptor affinities and the emotionally flavored terms exposed the compounds MDMA, MDA, LSD, 2C-C, and 25-I-NBOMe. The cortical brain regions expressing this complex of cell-surface receptor proteins preferentially occupied the extended parts of the visual cortex, but also in the superior parietal lobule and somatomotor cortex.

**Figure 2:**
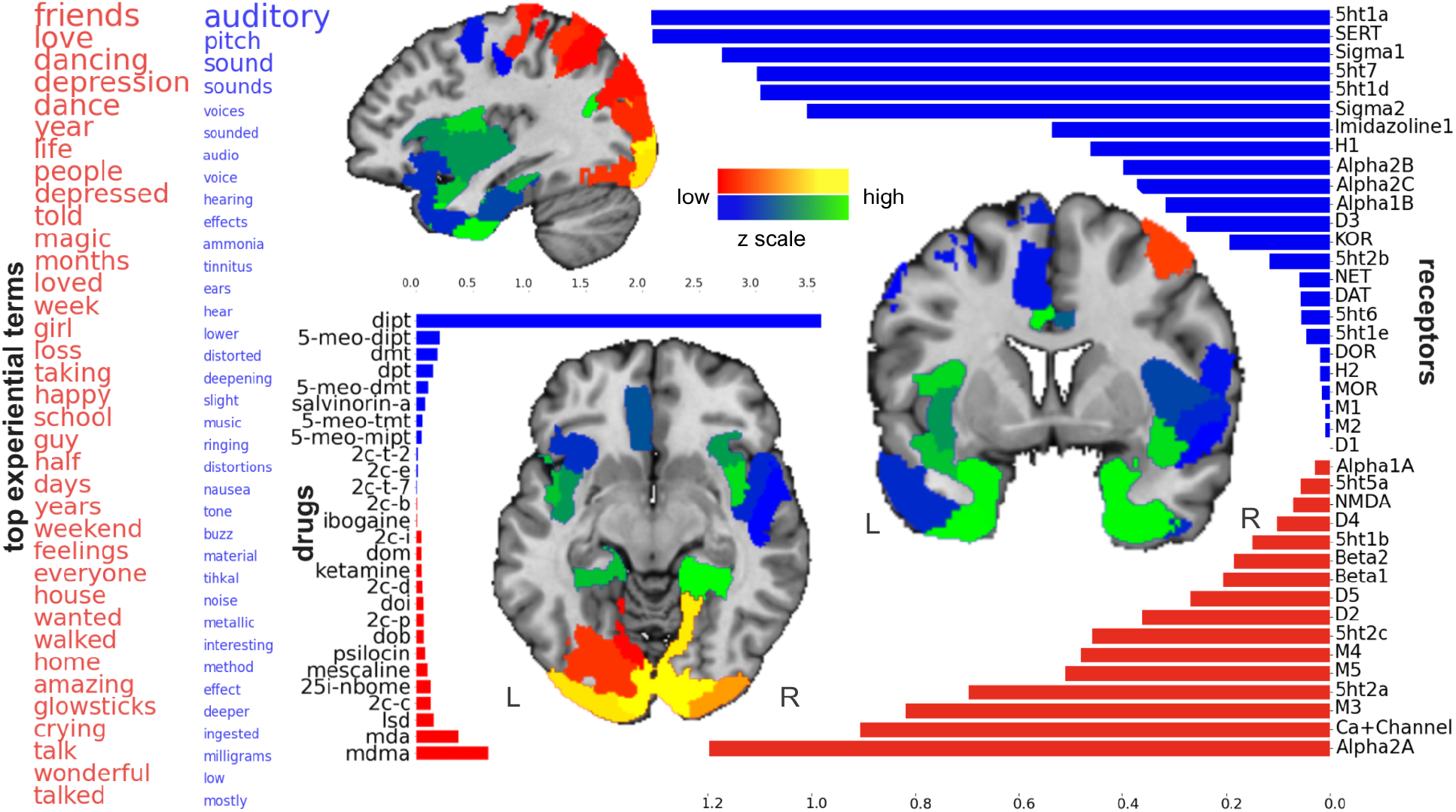
The second factor underlying hallucinogenic experiences highlights acoustic and emotional themes. Shows the second most important pattern of how binding affinities of neurotransmitter receptors were cross-linked with usages of >14,000 words in thousands of real-world hallucinogenic sessions. Sagittal, coronal, and axial brain slices are shown at x=-37, y=+4, and z=-9 (MNI reference space).

The opposite extreme of the second factor flagged a collection of terms with auditory sensation, including *auditory*, *audio*, *pitch*, *sound(s)*, *voice(s)*, *tone*, *ringing*, *music*, *buzz*, *hear*, and *hearing*, in addition to perturbations in sound perception like *distortion*, *deeper*, *tinnitus*, as well as the anatomical reference to *ears*. This experiential theme was linked to high affinity at molecular receptor 5-HT1A, in addition to lower-weighted affinity for SERT, Sigma 1, 5-HT7, 5-HT1D, and Sigma 2. These experience facets were also unique in being overwhelmingly associated with the DiPT drug. The auditory receptor-experience factor was preferentially localized to gene co-expression patterns in Heschl’s gyri, as well as the dorsomedial prefrontal cortex, dorsal anterior cingulate cortex, mid-cingulate cortex, medial temporal lobe extending into parahippocampal gyrus and temporal pole, and inferior parietal lobe. The primary auditory cortex in Heschl’s gyri are the first cortical regions to receive incoming auditory signals, while the DiPT drug was documented for causing characteristic distortions of auditory perception^30^.

In the third most important receptor-experience factor (explained variance *rho*=0.76, statistically significant at *p* < 0.05; Fig. 3), one extreme highlighted a theme of visual terms (*visual[s]*, *patterns*, *colors*, *eye*, *seeing*, *watch*), terms that suggest changes in normal vision (*movie*, *moving*, *intense*, *waves*, *tracers*, *swirling*), along with unpleasant physiological processes (*nausea*, *vomit*, *discomfort*, *tension*). These facets of experience were exclusively linked with molecular signaling via affinity to the serotonergic molecular receptors 5-HT2A, 5-HT2C, and 5-HT1A. This co-dependence of serotonergic affinities and experiential terms was linked to several phenethylamine and tryptamine drugs: DOI, 2C-T-7, 2C-D, 2C-C, 5-MeO-TMT, and 2C-P. The brain regions found to transcribe relevant combinations of synaptic receptor genes were expressed especially richly in the primary visual cortex, but also in adjacent visual processing regions including the posterior inferior temporal gyrus, fusiform facial area, and parts of the inferior parietal lobule.

**Figure 3:**
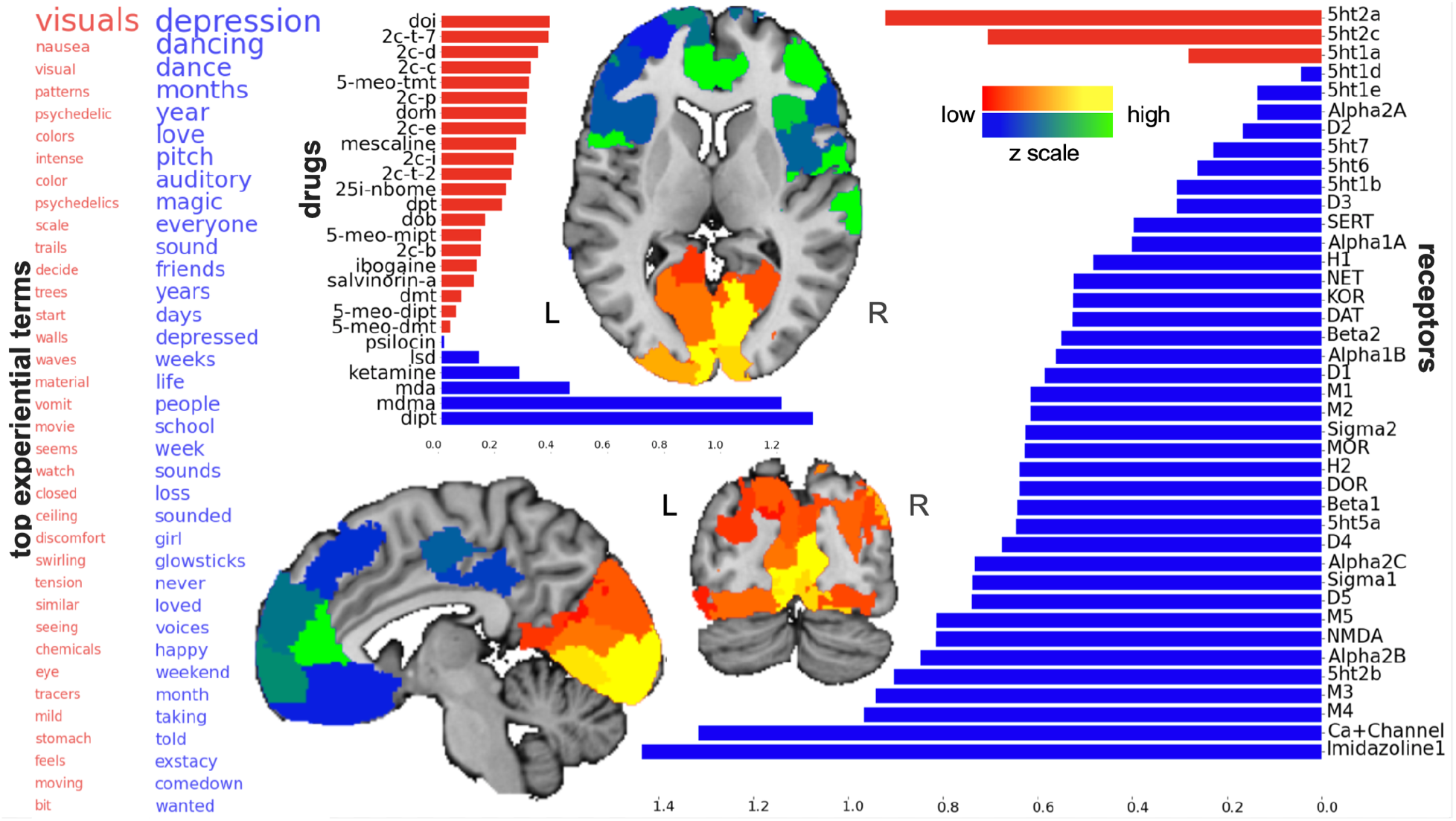
The third factor underlying hallucinogenic experiences highlights bodily and emotional themes. Shows the third most important pattern of how binding affinities of neurotransmitter receptors were cross-linked with usages of >14,000 words in thousands of real-world hallucinogenic sessions. Sagittal, coronal, and axial brain slices are shown atx=+6, y=-70, and z=+4 (MNI reference space).

The third factor spanned a range of experiential terms that was inhabited at the opposite extreme by emotional terms similar to factor 2, including *depression*, *depressed*, *dancing*, *love(d)*, *friends*, *loss*, *happy*, an intermediate time horizon suggested by the terms *days*, *weeks*, *months*, *years*, as well as some auditory terms. This constellation of experiential facets was preferentially linked to drugs with highest affinity for Imidazoline 1, calcium channel, M3, and NMDA receptors. This weighted receptor affinity was most closely associated with the drugs DiPT, MDMA, MDA, and ketamine. The anatomical brain regions that transcribe these combinations of cell-surface receptor proteins were highlighted in high-level association cortices such as the rostral anterior cingulate cortex, dorsolateral prefrontal cortex, temporoparietal junction, and bilateral insula.

The fourth most important receptor-experience factor (explained variance *rho*=0.71, statistically significant at *p* < 0.05; Supplementary Fig. 1) flagged term collections revolving around body load and somatic intensity (*euphoria*, *intense*, *breakthrough*, *energy*, *flame*, *loaded*, *scale*, *effects*, *dance*, *rush*) as well as several exteroceptive descriptors (*visuals*, *whispers*, *colors*). Similar to ego dissolution in factor 1, a theme of immediate time horizon was indicated by the term *seconds* and also implied by references to transient physiological parameters (*inhale*, *exhale*, *heart*). This constellation of experiential properties was linked most strongly to drug affinity with serotonergic (5-HT7, 5-HT1D, 5-HT2C) and adrenergic (Alpha 2c, Alpha 2b) molecular receptors. This pattern of receptor binding weightings was most closely associated with tryptamines DMT, 5-MeO-MIPT, and 5-MeO-DMT. The regions that transcribe this graded array of receptor proteins were limited to the right side of the brain and included most prominently the primary visual cortex, temporoparietal junction, posterior insula, and somewhat less intensely in the medial frontal gyrus and temporal pole.

At its opposite extreme, the fourth receptor-experience factor stressed terms relating to therapeutic treatment, including *plant*, *addiction*, *treatment*, *withdrawals*, and *methadone*. This factor also contained terms like *numb*, *dream*, *dizzy*, and *film* that may constitute a complementary theme of dissociation. Anatomical references to *bladder*, *leg*, *and feet* and the temporal modifier *years* were also associated with this receptor-experience factor. These cohesive constellations of conscious awareness were linked most strongly with drugs that preferentially bind NMDA, KOR, Sigma 1, D5, and D3 receptors. The drugs sharing the strongest affinity and experiential similarity with this receptor-experience constellation were ketamine, ibogaine, mescaline, salvinorin A, psilocin, and LSD. Brain regions found to preferentially transcribe combinations of these neurotransmitter receptor densities were located in the inferior parietal lobule, primary visual cortex, and extended areas of the premotor, motor, and somatosensory cortices.

The fifth most relevant factor (explained variance *rho*=0.68, statistically significant at *p* < 0.05; Supplementary Fig. 2) flagged the terms characterized by levels of comfort (*nausea*, *pain*, *numb*, *mild*, *burn*, *feels*, *pleasant*, *tolerance*) and other somatic terms (*nose*, *nostril*, *heart*, *body*, *feels*, *breathing*). The experiential horizon was temporally circumscribed by *minutes*. This constellation of conscious awareness alterations was strongly linked with drugs that preferentially bind to 5-HT1A, Sigma 1, and NMDA receptors. This fingerprint of receptor bindings was most especially associated with the drugs ketamine, 5-MeO-DMT, 2C-C, and DPT. The anatomical brain regions that transcribe these combinations of receptor genes preferentially in the primary visual cortex, dorsomedial prefrontal cortex, temporoparietal junction, and premotor cortex.

The fifth factor’s opposite extreme spanned a continuum of environmental terms (*plant, tree[s], green, sun, shapes, rain, sky, forest, life*). This theme was complemented by perceptual terms (*looked*, *saw*, *shapes*, *beautiful*, *whispers*, *laughing*, *images*) and references to *entities*, *beings*, and *faces*. This theme of subjective awareness alterations was linked preferentially with drug affinity at 5-HT2B, 5-HT5A, 5-HT1E, and 5-HT2C receptors as well as D1 and KOR. The derived profile of receptor implications was most closely associated with the drugs ibogaine, mescaline, LSD, and psilocin. The anatomical brain regions that transcribe these combinations of synaptic receptor genes located to the visual cortex and portions of the medial temporal lobe.

The sixth most relevant factor (explained variance *rho*=0.64, statistically significant at *p* < 0.05; Supplementary Fig. 3) highlighted a theme of euphoric sensations (*euphoria*, *high*, *nice*, *relaxed*, *enhanced*, *buzz*), perceptual impressions (*whispers*, *shapes*, *colors*, *geometric*, *patterns*, *music*), and consistent references to *entities* and *aliens*. This theme of alterations in subjective awareness was linked most strongly with drug affinity at serotonergic (5-HT2A, 5-HT2B, 5-HT2C), adrenergic (Alpha 2a, Alpha 2b), as well as NMDA and Sigma 1 receptors. The derived profile of receptor implications was mostly associated with the drugs 2C-C, DMT, 2C-D, ketamine, and 5-MeO-MIPT. The anatomical brain regions that co-expressed these receptor genes were in extended portions of the primary visual cortex, lateral prefrontal cortex, the temporoparietal junction, and the anterior insula.

At its opposite extreme, the sixth factor was characterized by intense fear (*terror*, *horrible*, *death*, *fear*, *panic*) and other dysphoric terms (*pressure*, *shit*, *throwing*, *horrible*, *dying*, *insane*, *surrender*). This experiential constellation also appears to contain a parallel theme of apparent transcendence (*forever*, *lightbulb*, *bliss*, *void*, *life*). This constellation of alterations of conscious awareness was linked with drugs that preferentially bind to serotonin receptors (5-HT1A, 5-HT7, 5-HT6, 5-HT1B) as well as D3 and D1. This fingerprint of receptor bindings was most strongly associated with the drugs 5-MeO-DMT, ibogaine, 5-MeO-TMT, and mescaline. The anatomical brain regions that transcribe these combinations of receptor gene expressions were in the medial inferior temporal lobe, inferior parietal lobule, and portions of the primary somatosensory and premotor cortices.

The seventh most relevant receptor-experience factor (explained variance *rho*=0.62, statistically significant at *p* < 0.05; Supplementary Fig. 4) emphasized terms relating to physical context, including *room*, *house*, *park*, *sat*, *walls*, *tent*, *street*, *outside*, and *school*. Apparent references to socio-emotional context were flagged by the terms *friends*, *mom*, *scared*, *panic*, *laughing*, *funny*, as well as *remember*, *talking*, *said*, *knew* and *thought*. This constellation of conscious awareness changes was induced especially by drugs that bind serotonergic receptors 5-HT1B, 5-HT1D, 5-HT7, 5-HT2A, 5-HT2C, 5-HT6 along with the D3 receptor. The drugs sharing the strongest affinity and experiential similarity with this receptor-experience constellation were 25-I-NBOMe, LSD, DOB, and psilocin. The brain regions that transcribe this graded array of receptor proteins occupied extended areas of the visual cortex, inferior temporal gyrus, a limited area of the inferior parietal lobule, bilateral temporoparietal junction, and portions of the somatosensory and motor cortices.

At its opposite extreme, the seventh receptor-experience factor flagged a collection of terms similar to factor 4 with references to therapeutic processes (*treatment*, *plant*, *addict*, *addiction*, *withdrawals*, *detox*, *provider*, *methadone*, *opiates*) as well as perceptual and somatic modifiers (*nausea*, *sick*, *stomach*, *ingested*, *visions*, *headache*, *pain*, *effects*, *stimulation*, *mild*, *pleasant*, *nice*, *relaxed*). This constellation of experiential features was linked most strongly to drug affinity with Sigma 2, NMDA, Imidazoline 1, DAT, and MOR molecular receptors. This pattern of receptor binding was overwhelmingly associated with the drug ibogaine. Brain regions found to preferentially transcribe combinations of these neurotransmitter receptor densities were located in the right dorsolateral prefrontal cortex, rostral anterior cingulate cortex, the bilateral anterior insula, and right temporoparietal junction.

The eighth and last factor revealed by our pattern-learning algorithm tracked joint variation between receptor affinity profiles and experiential terms (explained variance *rho*=0.60, statistically significant at *p* < 0.05). This factor delineated a theme of somatic relaxation, such as by flagging the terms *euphoria*, *relaxation*, *pleasant*, *enhanced*, *positive*, *impressed*, *stimulation*, *tactile*, and *waves*, along with contrasting terms such as *negative*, *anxiety*, and *terror*. These facets of drug-induced experience were associated with the timescale of *years*. The underlying emotional pattern was linked to a graded affinity for a narrow group of receptors: 5ht1a, 5ht1b, and D3. The cohesive variation between these receptor affinities and the emotionally flavored terms was linked to the drugs 2c-c, 2c-d, 5-meo-mipt, and salvinorin a. The cortical brain regions expressing this complex of cell-surface receptor proteins preferentially occupied the wider visual cortex, and portions of the superior parietal lobule and temporoparietal junction.

At its opposite end, the final factor flagged a collection of terms centered on emesis, including *vomiting*, *vomited*, *vomit*, *cramps*, *puke*, *nausea*, *bathroom*, and *shit*, in addition to terms of violence and force including *throwing*, *fucked*, *damage*, *and kick* as well as the anatomical reference to *eyes*. This experiential theme was linked most prominently to affinity at the Sigma 1 and NMDA receptors, in addition to a wide range of adrenergic and muscarinic receptors. These experience facets were strongly associated with the drugs 5-meo-tmt as well as DOI, DMT, and ketamine. This receptor-experience factor was preferentially localized to gene co-expression patterns in the anterior insula, dorsolateral prefrontal cortex, ventromedial prefrontal cortex, rostral anterior cingulate cortex, and orbitofrontal cortex.

## METHODS

### Rationale and workflow summary

Existing brain-imaging efforts to study psychedelics have typically concentrated on gross differences in neural activity after administering one particular drug. General principles of drug-induced brain changes cannot be derived by studying an isolated neurotransmitter system or a single drug. To overcome several of these limitations, our analytical strategy sought to chart principled patterns that link free-form testimonials of drug-induced experience with synaptic binding profiles of hallucinogenic drugs.

On the one hand, each text report was modeled as a histogram of word usages (‘bag-of-words’; e.g., one report may include the word ‘universe’ twice, ‘voices’ three times, ‘love’ once). This encoding of first-person narratives directly captured how individuals have articulated the changes in conscious awareness, which touches on thinking, perception, emotions, and other psychological alterations. On the other hand, each examined drug has documented binding strengths for specific receptor systems. For example, it is well known that LSD acts as a potent 5-HT2A agonist. Yet, this drug also has high affinity for a range of other serotonin receptors, in addition to exerting neuromodulatory effects on D3, D2, Alpha, and Beta receptors. Further, ketamine is typically described as an NMDA antagonist. However, this drug is also known to serve neuromodulatory roles on serotonin receptors as well as D2, DAT, SERT, and NET.

By invoking a doubly multivariate pattern-learning algorithm (canonical correlation analysis), we aimed to deconvolve cross-connections between constellations of neurotransmitter modulations and constellations of experiential terms, which cut across 6,850 written reports of psychedelic experience. In this bottom-up approach, we have also explicitly modeled the possible co-existence of several separate receptor-experience factors. That is, each unique report of hallucinogenic experience was modeled as a weighted combination of distinct receptor-experience factors to be discovered. In an attempt to complement previous psychedelics research, we thus designed an analytical framework that can extract several hidden representations of rich subjective experiences across a diversity of hallucinogenic drugs.

### Experience samples of psychedelic drug experiences from Erowid resource

Erowid Center is a not-for-profit, member-supported organization. This initiative hosts a collection of first-hand accounts of experiences that were elicited by psychoactive drugs. The library aims to educate the public about the expected and unwanted consequences of each drug to help fight misuse or harm. This web platform (www.erowid.org) aims to provide an educational resource on the effects of psychoactive substances – both legal and illegal ones. This overall collection now includes >38,000 testimonials in English language in free-text form from >800 different drugs, plants, and fungi. The testimonials have been written and submitted without any financial incentives. There were no constraints on the length, format, and no externally imposed structure on the content of the reports. These real-world experiences of psychoactives provided a unique window into the effects of these compounds. The database of drug narratives from Erowid’s ‘experience vault’ is entirely anonymous.

To pursue the goal of the present study, we derived a dataset that comprised a total of 6,850 experience reports. Each report belonged to one of the 27 drugs. This final dataset was obtained based on the following eligibility criteria for inclusion of experience reports:

1. Compounds with a high number of testimonials in the Erowid database.
2. Compounds with well-known receptor binding affinity^22,31^.
3. Compounds that are believed to mediate effects through multiple receptor systems.
4. Classical psychedelics.
5. Closely-related research chemicals purported to have psychedelic-like effects, including substituted phenethylamines, tryptamines.
6. Drugs known to induce profound reconfigurations of conscious awareness^1^ in particular ego-dissolution: classical psychedelics, entactogens, dissociative anesthetics, and agonists of the kappa opioid receptor (KOR)^32^.

### Natural language processing pipeline

To enable receptor-experience modeling, we initially constructed a bag-of-words encoding of the text descriptions in each testimonial. To build this vocabulary representation, we started by removing punctuation marks and special characters. Next, we discarded words that consisted of only 1 or 2 characters. We also discarded words that occurred less than 7 times in the entire corpus of experience reports. We then removed common prepositions, pronouns, drug names and articles. Subsequently, all words in the dictionary were lower-cased for the sake of consistency. Using the ensuing dictionary, we tabulated the counts of each word for each testimonial. This word encoding tactic yielded a summary matrix with testimonials in the rows and 14,410 unique words indexed in the columns. This word matrix *M* was sparsely populated because most words do not occur in most testimonials.

To preprocess the testimonial-word matrix *M*, we have used the *term-frequency inverse-document-frequency* (tf-idf) transformation that is widely used for text mining in the natural language processing community^33^. The metric allowed us to compute a notion of word frequency in a given testimonial in a way that carefully accounts for its global prevalence in our entire corpus. The tf-idf values for a given word increased by how informative each particular word was within a testimonial (tf) and decreased with the frequency with which the word was used ubiquitously across the whole corpus of testimonials (idf). The term frequency (tf) is commonly interpreted as the similarity of the effects that the words have in a given testimonial in which they occur. The inverse document frequency (idf) is defined as the logarithmically scaled inverse fraction of the documents containing the term. We thus used a term frequency adjusted for document length and smooth inverse document frequencies by computing the product of the two statistical aggregate quantities:

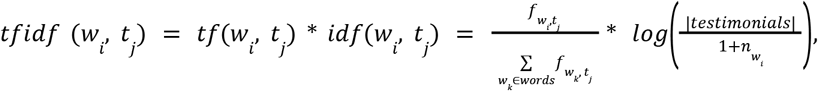

where *w_i_*, denotes the word vocabulary entry in column *i*, *t_j_*. refers to the testimonial in row *j*, *f_w,t_* is the frequency of word *w* in testimonial *t*. Hence, 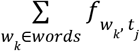 computes the total number of words in testimonial*t_j_* and *n_w_* is the count of testimonials containing word *w*. In this way, a higher tf-idf value directly reflected the salience of a used word for the meaning of a particular testimonial, but was simultaneously discounted as a function of abundance in the corpus in general. One benefit of this encoding scheme is effective handling of very common words in the corpus.

Each row in the tf-idf transformed word count matrix *M* encapsulated the rich semantic aspects of one lived experience with a hallucinogenic drug. Counts from words that tend to co-occur in the same documents or words that co-occur in the same contexts created correlations along the rows and columns of this matrix. This type of bag-of-words representation is naive about word order and thus ignored event sequences in the experience reports. Nevertheless, this word encoding scheme is known to capture high-granularity semantic information in many text-mining applications^33^.

To automatically search and organize the space of semantic representations in the experience reports, latent semantic analysis (LSA) was a natural choice of pattern-learning algorithm^34^. Applying LSA to the corpus of preprocessed testimonials, we extracted a set of obtained semantic components of contextual word usage, which were ordered from most to least important based on the explained variance in summarizing word usage combinations. We extracted the top *k*=800 semantic components from the word count matrix *M* for further analysis (cf. below). Each testimonial was thus decomposed into a vector of *k* component projections whose value is the sum of vectors standing for its component words.

More formally, LSA extracted semantic representations from the sparse rectangular testimonial-word matrix *M* by decomposition into three separate matrices:

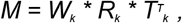

where the matrix *W*_|words|x k_ carries orthonormal columns representing word weights, the diagonal matrix *R*_k x k_ carries component relevances ordered from most to least explanatory, and the matrix *T*_|testimonials| x k_ carries orthonormal columns representing testimonial weights that together spanned out the semantic space of the examined corpus of hallucinogenic experience reports.

In doing so, LSA detects and extracts the similarity of collective words that co-occur in a testimonial to the extent to which they are attributable to a common semantic context. Conversely, LSA can be viewed as modeling the approximate meaning of testimonials as the average of the meaning of their words. It has been conjectured that, in many applications of LSA, the optimal dimensionality is intrinsic to the analyzed domain and must be therefore empirically determined. In the present investigation, we have benefitted from canonical correlation analysis to explore joint modeling solutions constrained by drug receptor affinity matrix to find an optimal degree of data reduction (cf. next passage).

To counteract class imbalance between the available testimonials per drug, we have drawn a random sample of 300 testimonials from each particular drug. Sensitivity analyses have confirmed that similar semantic components are found when applying LSA to different subsets of testimonials. Different sampling proportions of drug testimonials have changed the ordering of the components, but the underlying semantic components themselves proved to be robust. Further, we have committed to *k*=800 semantic components, preserving 60.7% of the variation in the corpus testimonials, for all subsequent analyses. Sensitivity analyses here showed that varying the number of LSA components between *k*=50-5000 components, preserving 50.0%-99.9% of the variance in the testimonial-word matrices *M*, gave substantially similar results and led to the same neuroscientific conclusions. This setup resulted in a final vocabulary of 14,410 words in the dictionary.

### Constructing receptor affinity fingerprints

In order to elucidate the spatial distribution of hallucinogenic compounds that modulate neuronal activity throughout the cortex during subjective “trips”, we compared normalized measurements of their relative binding strengths. For each of the 27 drugs to be analyzed, we built an affinity vector that captures the binding strengths (Ki) for 40 targets: G protein-coupled receptors (GPCRs), molecular transporters, and ion channels. Binding data was aggregated primarily from two sources: (Ray 2010)^22^ and Rickli (2015)^31^, while values from ketamine were found in PDSP database. In the cited studies, of all sites tested only at 40 was significant affinity measured. These sites included 5-HT receptors (5-HT2A, 5-HT2C, 5-HT2B, 5-HT1A, 5-HT1B, 5-HT1D, 5-HT1E, 5-HT5A, 5-HT6, 5-HT7), dopamine receptors (D1, D2, D3, D4, D5), adrenergic receptors (*α*1A, *α*1B, *α*2A, *α*2B, *α*2C, *β*1, *β*2), serotonin transporter (SERT), DA transporter (DAT), norepinephrine transporter (NET), imidazoline1 receptor (I1), Sigma receptors (*σ*1, *σ*2), delta opioid receptor (DOR), kappa opioid receptor (KOR), mu opioid receptor (MOR), muscarinic receptors (M1, M2, M3, M4, M5), histamine receptors (H1, H2), calcium ion channel (Ca+) and N-methyl D-aspartate (NMDA) glutamate receptor.

The binding values for 20 of the hallucinogenic drugs found in Ray (2010) were from new (16 compounds) or existing (LSD, salvinorin A, 5-MeO-TMT) binding assays performed in coordination with the NIMH Psychoactive Drug Screening Program (PDSP). The methodology of the new binding studies followed Glennon, et al^35^: briefly, for each compound, a primary assay at 10nM concentration was performed against each receptor, transporter, or ion channel. Those compounds that induced a “hit” of >50% inhibition were then subjected to a secondary assay at 1, 10, 100, 1,000, and 10,000 nM to determine Ki values, with the final value calculated as the average of at least three replicated assays. Further details of how individual assays were conducted can be found at (https://pdsp.unc.edu/databases/kidb.php). The binding data found in Ray (2010) for ibogaine was collected from the literature^22^. The binding data for 6 additional drugs (25-I-NBOMe, 2C-C, 2C-D, 2C-P, 2C-I, 2C-T-7) were collected from studies performed by researchers at the University of Basel in Switzerland^31^, the methodology for which has been explored in detail elsewhere^36^. Finally, the affinity vector for ketamine was calculated from additional studies contained in the PDSP database, and presented as the average of all Ki values found (https://pdsp.unc.edu/databases/kidb.php). In addition to NMDA ketamine was found to have significant binding values for Sigma opioid receptors, an outdated nomenclature which is now differentiated into Sigma 1 and Sigma 2 receptors. A sensitivity analysis was performed and we found that utilizing both values as negative, positive for Sigma 1 only, and positive for Sigma 1 and 2 all yielded indistinguishable final results. The sources for all studies cited can be found in supplementary material (STable 1 and 2).

To enable apples-to-apples comparisons between different drugs in terms of their spatial concentration within cortical tissue, our study relied on the inhibitory constant (*K*_I_): a raw measure of the molar concentration (i.e., absolute value) that is necessary to displace 50% of a known ligand from the binding site of a given cell-surface receptor. While such Ki values do not capture differences in functional activity induced by specific drugs at these targets, our ultimate aim was instead to contrast where these compounds act in the brain. Ki values are an optimal proxy for this purpose of simulating spatial distributions of drug-receptor binding during hallucinogenic experiences. This standard measurement of receptor affinity in pharmacology is computed as follows:

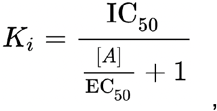

where *[A]* denotes the fixed concentration of agonist and *EC*_50_ is the excitatory concentration of agonist that results in half maximal stimulation of the receptor. The *IC*_50_ value for a drug is the inhibitory concentration to reduce the biological activity to half of its maximal activity on a receptor. The resulting *K_I_* is the inhibition constant for a drug: the concentration of a ligand in a competition assay which would occupy 50% of the receptors if no ligand were present.

Since higher binding affinities are represented by lower *K_I_* values that span orders of magnitude ^22,37^, a negative log10 was performed. In order to control for the confounding factor of drug potency, we followed the procedure of Ray (2010): npKi = 4+pKi - pKiMax. In instances where there was no available binding data for a given receptor or the binding was nonsignificant (>10,000nm), the value was set to the minimum value. The advantages of these normalization procedures are that they result in higher affinities having higher numerical values, affinities that are very low are reported as zero, the highest affinity is set to 4, each unit of npKi is reported as one order of magnitude of the crude Ki value, and potencies are factored out so that drugs of varying potency can be directly compared in terms of their affinity.

A virtual screening of binding affinity profiles for the drugs described by Ray (2010) was previously performed^21^ using the method introduced by Vidal and Maestros (2010). This procedure found that the binding strength between a ligand and a particular target can be approximated based on its structural resemblance to a selection of previously assayed compounds. The molecular structure of each compound was described as a binary vector, and the structural similarity between compounds was calculated using the Tanimoto distance coefficient. The Ki values for each drug at 34 receptors were then predicted in terms of their structural similarity and compared against the known binding affinity of 13 non-selective psychiatric drugs. This analysis found 39% of hits against 34 targets, which was considered an acceptably positive result compared to the reference value of 59-68% found by Vidal and Mestres (2010) using the same procedure on a larger database of compounds.

### Exploring coherent receptor-experience patterns underlying psychedelic episodes

Next, we sought dominant factors – “modes” of joint variation – that provide insight into how semantic components emerging from word usage patterns are inter-linked with receptor binding affinities from 40 neurotransmitter receptor subclasses. The canonical correlation analysis (CCA) algorithm was ideally suited to interrogate the possible existence of such a multi-modal correspondence between two high-dimensional variable sets^38,39^. A first variable set *X*_|testimonials| x k_ was constructed from LSA-derived re-representation of experience reports by means of the discovered semantic components (cf. above). A second variable set *Y*_|testimonials| x 40_ was constructed from each drug’s known pharmacological properties regarding molecular affinity to the neurotransmitter receptors. The affinity profile vector of each drug corresponded to the drug taken during each testimonial. Unmeasurable affinity values and values for which there was no available data were imputed with the minimum affinity level for the purpose of the present investigation. CCA involved computation of the projection vectors *a* and *b* that maximize the relationship between a linear combination of semantic contexts (*X*) and a linear combination of receptor affinity profiles (*Y*) across testimonials. CCA thus searched a large space of possible combinations by identifying the two projections *Xa* and *Yb* that yielded maximal association between the semantic context features of the drug experience and the neurotransmitters in the brain to which the drug binds.

More formally, we have solved the doubly multivariate matrix decomposition problem based on the following optimization objective:

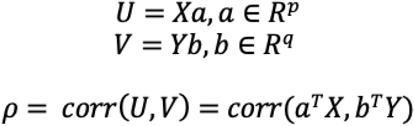

In addition to maximizing the correspondence between the variable sets *X* and *Y* as the first canonical factor *h*, it is possible to continue to seek additional pairs of linear combinations that are uncorrelated with the first canonical mode(s). This process may be continued up to *min*(*p*, *q*) times. While *a* and *b* here denote the *canonical vectors*, *U* and *V* are commonly referred to as the *canonical variates*. The *canonical correlation* is a performance statistic that denotes the *correlation correlation* rho between the canonical variates for each factor *h*.

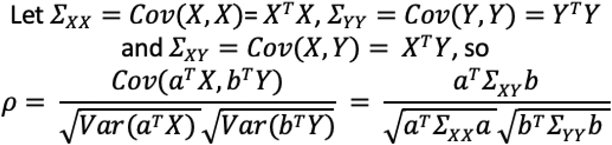

The relationship between our original data (*X* and *Y*) and the derived canonical variates (*U*_|testimonials| x h_ and *V*_|testimonials| x h_) can be understood as the best way to rotate the left variable set and the right variable set from their original representation to new representations that maximize their linear association. The fitted parameter values of the CCA model thus described the rotation of the coordinate systems: the canonical vectors encapsulating how to transition from the original measurement coordinate system to the new hidden space, the canonical variates encapsulating the position of each data point (i.e., testimonial or drug experience) in that new embedding space.

The set of *h* mutually uncorrelated (i.e., orthogonal) factors of joint variation were modeled with the optimization constraint to be linearly independent from each other. CCA also yielded factors that were already ordered from the most to least important receptor-experience pattern. The first and strongest factor therefore explained the largest fraction of covariance between molecular features of receptor pharmacology and experiential features of semantic context. Each particular factor captured a unique fraction of the multivariate co-variation in the Erowid corpus that was not explained by one of the *h* - 1 other factors.

To assess statistical significance, we have determined the robustness of each estimated CCA factor by hypothesis testing in a non-parametric permutation procedure used previously^38,40^. Relying on minimal modeling assumptions, a valid null distribution was derived for the correlation between canonical variates that resulted from CCA analysis. In 1,000 permutation iterations, the semantic context matrix *X* was held constant, while the receptor affinity matrix *Y* was subject to testimonial-wise random shuffling. The constructed surrogate data preserved the statistical structure idiosyncratic to the type of our input data, yet permitted to selectively destroy the signal properties related to the CCA statistics to be tested^41^. The thus generated empirical distribution reflected the null hypothesis of random association between semantic context features and pharmacological drug features across testimonials. *P* values were obtained from the number of canonical correlations rho that exceeded the null CCA model of maximum correlation values. This approach yielded *h*=8 significant CCA factors of receptor-experience correspondence, where explicit correction for multiple comparisons was carried out searching through all estimated CCA factors (all *p* < 0.05, family-wise-error-corrected). Each explained-variance metric rho indicated a given factor’s strength of variation that is common to both semantic contexts of experience reports and drug-specific receptor binding profiles.

### Mapping receptor-experience factors to regional levels of receptor gene transcription

We finally sought to anatomically locate the significant receptor-experience factors based on their gene transcription weighting of neurotransmitter receptor subclasses. Publicly available human gene expression data from six whole postmortem brains of neurotypical donors (one female, aged 24 years to 57 years [42.5 ± 13.38 years]) were obtained from the Allen Human Brain Atlas (http://human.brain-map.org). To strengthen reproducibility and comparability, we have used the abagen tool to map the receptor gene expression information to the 200 Schaefer-Yeo regions (https://github.com/rmarkello/abagen). Averages of invasive brain tissue probes were computed across all six donors for each of the 40 receptor systems of interest.

For each of the 200 anatomical brain regions *z*, we have then derived the implication of a given receptor-experience factor *h* by computing the following dot product:

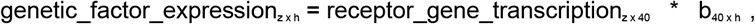

where *receptor_gene_transcription* denotes the (z-scored) measure of gene expression activity for a specific brain region *z. b* denotes the canonical vector of a particular factor *h* that carries the linear combination obtained from fitting receptor binding affinity profiles (*Y*) from the CCA step of our analysis (cf. above). The transcriptomic expression data were transformed to a minimum gene transcript expression of zero to ensure the directionality of the CCA factors. The ensuing region-wise quantities indicated the degree to which the semantic structure captured by LSA is linked to variation in the receptor expression profile of a specific brain region. The ensuing region-specific estimates of genetic factor expression were demeaned and variance-scaled for the sake of visualization. The factor-specific co-expression of receptor genes was directly topographically mapped to each of the 200 target brain regions of the Schaefer-Yeo reference atlas.

More generally, the DNA microarray probes of post-mortem brain tissue have previously been successfully used for genome-wide mapping to build transcriptomic cortical atlases^42^. The regional levels of gene transcription have been reported to closely track the regional levels of actual protein expression across the cortex^43^. Indeed, in-vivo positron-emission tomography imaging in humans was also closely aligned with the spatial gene expression levels from this dataset^44^. For these reasons, the high-throughput transcriptomic data from the Allen Institute, leveraged in the present study, was shown before to allow rigorous mapping of interregional variation in proxies of synaptic receptor densities.

### Scientific computing implementation

Python was selected as the scientific computing engine. Capitalizing on its open-source ecosystem helps enhance replicability, reusability, and provenance tracking. The scikit-learn package provided efficient, unit-tested implementations of state-of-the-art machine learning algorithms (http://scikit-learn.org). This general-purpose machine-learning library was interfaced with the nilearn library for design and efficient execution of brain-imaging data analysis workflows (http://github.com/nilearn/nileam). Data plots were generated by Matplotlib (https://github.com/matplotlib/matplotlib).

All scripts for the analysis pipelines that reproduce the results of the present study are readily accessible and open for reuse by the reader: https://github.com/lucidtronix/Trips-and-Neurotransmitters-Discovering-Principled-Patterns-across-6850-Hallucinogenic-Experiences

## DISCUSSION

Subjective psychedelic experience is commonly studied in the strictly controlled laboratory setting: after participants consume a particular hallucinogenic drug and lie down in the bore of a brain-imaging scanner, their functional activity changes are measured and compared to those of control subjects. Our study confronts the neurophysiological basis of hallucinogenic states of consciousness from a different direction: we have translated natural language processing tools from machine learning to mine a large corpus of 6,850 real-world narratives with 27 different drugs. By adopting a data-driven strategy, we sought to uncover general principles that delineate how drug-induced changes in subjective conscious awareness are anatomically rooted in 40 neurotransmitter receptors. Cutting across a variety of drugs, we have assigned concrete numbers to distinct facets of psychedelic episodes and have mapped differences in transcriptomic proxies of synaptic receptor densities onto cortical regions. In so doing, we have uncovered a set of complementary principles that describe how different aspects of subjective experience may be triggered by receptor modulation in specific cortical regions. We have developed a framework that can inform the design of new classification systems and treatment options for psychoactive substances in the future.

To deepen understanding of the neurobiological and behavioral effects of hallucinogenic drugs, it is beneficial, we argue, to consider its action within defined neurotransmitter systems and their transcriptomic gene expression within different anatomical regions. Since the 1990’s, it is known that predosing with 5-HT2A-antagonist ketanserin can negate the effects of psilocybin in a dose dependent manner^45^. Similarly, the effects of KOR-agonist salvinorin A and NMDA-antagonist ketamine have been found to be attenuated by predosing with specific substances^46 47^. However, ibogaine is known to have strong binding affinity at all of these cell surface receptors. In fact, ibogaine’s action at 5-HT2A, KOR, and NMDA receptors^48^ leads to a unique profile of psychological changes^49^. These receptor-based studies have underscored that certain key receptors are necessary to mediate specific psychoactive properties. Yet, this handful of receptors are unlikely to be a complete description of what explains the inter-individual variations and nuances described in hallucinogenic experiences. As the most important result of our investigation, we offer a framework for understanding how constellations of neurotransmitter pathways – rather than single-receptor actions – may contribute to psychedelic experiences.

Ego dissolution is a hallmark of hallucinogenic states of consciousness. This experience frequently involves blurred self-world boundaries, oceanic boundlessness, and out-of-body illusions. Earlier research on ego dissolution has placed a focus on binding to the 5-HT2A receptor^50–52^. However, clinical surveys using standardized instruments for ego dissolution reveal that other classes of drugs also consistently produce such experiences, including the KOR-agonist salvinorin A^24,53^ and NMDA-antagonist ketamine^23^. In line with these earlier findings, our analyses have established close links between disrupted self-experience and drug affinity to KOR, D1, 5-HT7, and NMDA receptors. Indeed, salvinorin A, LSD, and other drugs bind the D2 receptor, which is the primary target of antipsychotic drug treatment. This receptor subclass stimulation may contribute to the loosening of boundaries between self and environment, which is a central feature of psychosis^54^. In addition, KOR, 5-HT7, and NMDA receptors are targets of novel antipsychotic drugs. Taken together, a wider group of receptors may individually or collectively underpin the process of self-disintegration that is thought to be critical for the success of psychedelics-assisted psychotherapy^17–19^.

Previous clinical assessment tools have been designed to accurately detect ego dissolution^6,7,55^ and identify corresponding changes in functional network connectivity in the brain^8,50^. Our study locates these subjective states within cortex-wide patterns of neurotransmitter receptor gene expressions. Sieving through a rich vocabulary of >14,000 words, we found that the leading receptor-experience factor tracks a theme of mental expansion, indicated by the terms *consciousness*, *earth*, *reality*, *existence*, *space*, and *universe*. The top factor also revealed references to liminal beings through the terms *alien*, *entity*, *sitter*, *spirit*, and *beings*. This constellation of experiential descriptions resonates with various measures of the mystical experience^6^. Indeed, a recent questionnaire study that reported the phrase “I felt at one with the universe” to be intimately related to the experience of ego dissolution^55^.

This experience of self-disintegration has been argued to be linked to weakening of the default mode network’s organizing influence on neural activity in subordinate brain networks^28,29^. Loosening the grip of top-down control exerted by the highest integrative network likely leads to leaky filtering of sensations, thoughts, and feelings^26,27^. Consistent with this view, factor 1 highlighted the inferior parietal lobule, retrosplenial cortex, and dorsal posterior cingulate cortex within the default mode network (DMN) in conjunction with the theme of ego dissolution. However, our study also points to a role of the salience network as another high-level center of integrative processing that was strongly highlighted in factor 1, which has been less often mentioned in early neuroimaging studies on psychedelics. In contrast, the opposing perceptual pole highlighted lower-level networks, such as the wider visual cortex. This polar distribution of receptor expression reflects the functional antagonism between the higher associative networks-overarching regulators of brain-wide neural activity – and subordinate network systems more closely linked to perception, emotion, and action. Simultaneous modulation of these neuronal circuits by KOR, 5-HT7, D2, NMDA, and 5-HT2A may ultimately underpin experiences of self-transcendence.

Invasive electrical stimulation of the inferior parietal default mode network has been found to alter own-body perception, and even lead to experiences of whole-body displacement^56^. Moreover, changes in within-DMN functional connectivity, especially in its inferior parietal regions, were shown to explain the most global variation in the functional interplay between major brain networks^57^. These changes in connectivity appear to mediate therapeutic effects, as psilocybin-induced changes in the DMN activity even predicted behavioral differences 4 weeks after drug consumption. Similarly, the salience network has been called ‘a link between the systems’^58^. This coinage is due to this network’s involvement in global monitoring and control of action-perception cycles to influence various executive functions^59^. Lesions in the mid-cingulate cortex or anterior insula of this canonical network indeed were reported to disrupt attentional reallocation and action initiation in structured task execution^60,61^.

Our analysis also uncovered hidden relationships between receptor affinities and perceptual themes that both confirm prior findings and suggest exciting new directions for future research. Visual distortions are ubiquitous in hallucinogenic states and are commonly characterized by vivid colors and fractal patterns. Indeed, *visuals* emerged as the top hit among >14,000 candidate terms in several significant receptor-experience factors in our study. Prior research has indicated that stimulation of 5-HT2A receptors concentrated in layer 5 pyramidal neurons can induce elementary (fractal) and more complex visualizations^25^. However, the affinity weightings of our receptor-experience factors suggest important roles for other brain regions in distorting visual perception. While the visual cortex was predictably most strongly linked to these visual changes, the medial temporal lobe, ventromedial prefrontal cortex, dorsolateral prefrontal cortex, and rostral anterior cingulate cortex were also consistently implicated in an array of serotonergic receptors. These findings implicate a variety of specific regions and receptors (5-HT7, 5-HT2C, 5-HT1A, Alpha 2a) that may play important roles in instantiating the powerful and diverse array of visual changes that hallucinogens can elicit.

Moreover, psychedelic trips frequently alter time perception or understanding of time. Undergoing ego dissolution is typically accompanied by a perceived breakdown of the space-time continuum^1,62^. Consistently, the leading factor was selectively linked to momentary aspects of hallucinogenic experience, such as *eyes* and *lungs*, along with fleeting physiological functions, including *inhaled*, *exhaled*, and *breath*. In our findings, this theme was specifically tuned to experience reports that contained the words *seconds* and *hours*. This short time horizon was not salient in the second and third most explanatory receptor-experience factor. Instead, these separate sources of brain-behavior variation were dominated by emotional themes and featured words of a longer time scale, including *days*, *weeks*, *months*, *year*, *weekend*, and *life*. Hence, the three chief factors of hallucinogenic states of consciousness have revealed strong and opposite relations to drug-induced changes of the internal clock.

Hallucinogenic states of consciousness have been described to stretch minutes to feel like hours or, conversely, to compress hours into seconds in subjective time perception^1,62^. Our approach may shed light on the array of neurotransmitter receptor bindings that underlie different types of altered experiences of time. In particular, the role of the dopamine receptor system in our leading factors appeared to co-occur with the immediate time horizon. D1 receptors were correlated with the immediate time horizon of seconds in factor 1 (mystical experience). Instead, D2, D4, and D5 were related to the longer timescales of weeks, months, and years in factors 2 and 3. Our algorithmic decomposition of experience reports appears to be in line with invasive experiments on time perception in animals. In mice, suppression of dopamine neurons decreased behavioral sensitivity to time^63^. Neural activity in dopamine neurons was demonstrated to track time estimates between experimental trials. Using drugs or optogenetic intervention on these dopaminergic neuron populations, mice made poorer decisions that involved internally keeping track of temporal information. Individuals with schizophrenia are also notoriously impaired in the processing of temporal information. Such patients have a generally distorted sense of time, with an overlying variable internal clock^64,65^. Patients carrying a diagnosis of schizophrenia are commonly treated by drugs that typically have dopamine-antagonism as an important component of their neurophysiological action^66,67^. Moreover, stimulation of the 5ht2 receptor has also been shown to mediate time-contingent action in rats^68^ and humans^69,70^.

In sum, distinct receptor-experience factors revealed distinct distortions of time perception. This key finding shows that a) several of our data-driven findings dovetail with experimental, invasive evidence in animals, b) the drug-induced ‘experience of time’ may actually affect more than the dopamine receptor system alone, and c) different types of ‘experience of time’ are potentially linked to different receptor binding constellations.

### Conclusion

Hallucinogenic drugs are opening a door into the primary biology that instantiates conscious awareness. These causal interventions on specific neuronal circuits are generally non-toxic, do not make users addicted, and directly affect only the brain. By innovating analytical strategies for data-fusion, we have coalesced three separate windows into cognition: 1) several thousand real-world anecdotes about hallucinogenic drug episodes, 2) synaptic binding affinities for various neurotransmitter systems, and 3) human brain tissue probes of receptor gene transcription. By bringing into contact these disparate levels of investigation, our study could detail the consequences of the chaotic breakdown of the hierarchical functional organization.

Changes in functional interaction among large-scale networks probably induce rebalancing between bottom-up sensory signaling and highly integrative top-down control. In all our leading explanatory factors, hallucinogenic consciousness states have pinpointed manipulation at both extremes of the neural processing hierarchy: the deepest layers of our neural networks, especially the higher-order association circuits, as well as the shallowest layers of brain hierarchy, especially the primary visual cortex. The des-integration of ordinary regimes of top-down influence may index the subjective confusion between internal sensations and facts of the external world.

Ultimately, psychedelics may come to serve as a tool to study the mechanistic receptor basis of how the higher association cortex orchestrates its subordinate large-scale circuits to impose structure on sensory perception. Hallucinogens have a meandering history. Yet, they offer potential to break the long drought in novel drug therapies for psychiatric patients.

## Supporting information

Receptor drug affinities

Gene name to receptor mappings

## ACKNOWLEDGEMENTS

We are grateful to Ross Markello for technical consulting on the neurogenetic analysis workflow using the abagen tool. We thank the founders, curators, contributors, and volunteers of Erowid Center for sharing their data and for their decades of work on the experience report collection.

This project has been made possible by the Brain Canada Foundation, through the Canada Brain Research Fund, as well as by NIH grant R01AG068563A and the Canadian Institutes of Health Research. DB was also supported by the Healthy Brains Healthy Lives initiative (Canada First Research Excellence fund), and by the CIFAR Artificial Intelligence Chairs program (Canada Institute for Advanced Research), as well as Research Award and Teaching Award by Google.

## AUTHOR CONTRIBUTIONS

G.B., S.F.F., and D.B. conceived the study, interpreted the results and wrote the manuscript. D.B. led data analysis.

## COMPETING INTEREST

The authors declare no conflict of interest.

## SUPPLEMENTARY ONLINE MATERIAL

**Supplementary Figure 1: The fourth factor underlying hallucinogenic experiences.** Sagittal, coronal, and axial brain slices are shown at x=20, y=-43, and z=8 (MNI reference space).

**Supplementary Figure 2: The fifth factor underlying hallucinogenic experiences.** Sagittal, coronal, and axial brain slices are shown at x=11, y=-15, and z=2 (MNI reference space).

**Supplementary Figure 3: The sixth factor underlying hallucinogenic experiences.** Sagittal, coronal, and axial brain slices are shown at x=5, 28, y=-26, and z=23 (MNI reference space).

**Supplementary Figure 4: The seventh factor underlying hallucinogenic experiences.** Sagittal, coronal, and axial brain slices are shown at x=7, y=-23, and z=7 (MNI reference space).

**Supplementary Figure 5: The eighth factor underlying hallucinogenic experiences.** Sagittal, coronal, and axial brain slices are shown at x=-6, y=-17, and z=6 (MNI reference space).

**Supplementary Table 1: Receptor affinity matrix**

**Supplementary Table 2: Molecular receptors and gene names in the Allen Human Brain Atlas**

## Notes

### Competing Interest Statement

The authors have declared no competing interest.

### Summary of Updates

Added more supplementary data

