## Supplementary Figures for "Trips and Neurotransmitters: Discovering Principled Patterns across 6,850 Hallucinogenic Experiences"

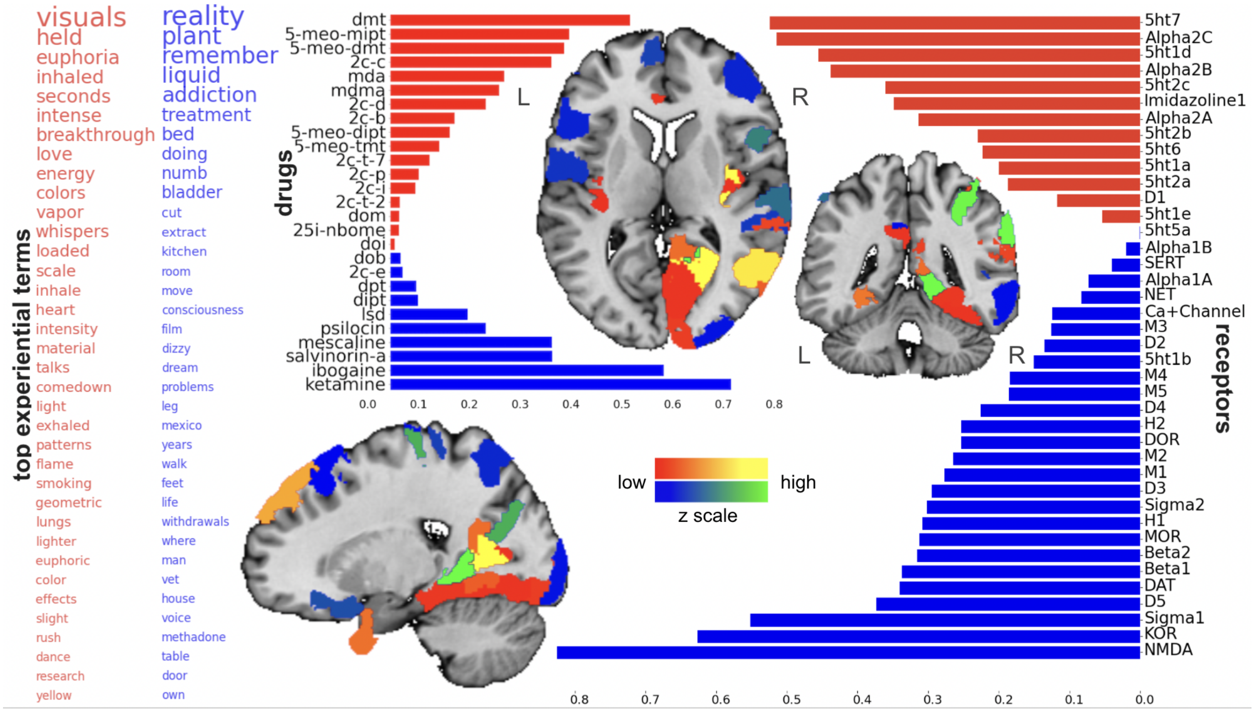


**Supplementary Figure 1: The fourth factor underlying hallucinogenic experiences.** Sagittal, coronal, and axial brain slices are shown at x=20, y=-43, and z=8 (MNI reference space).

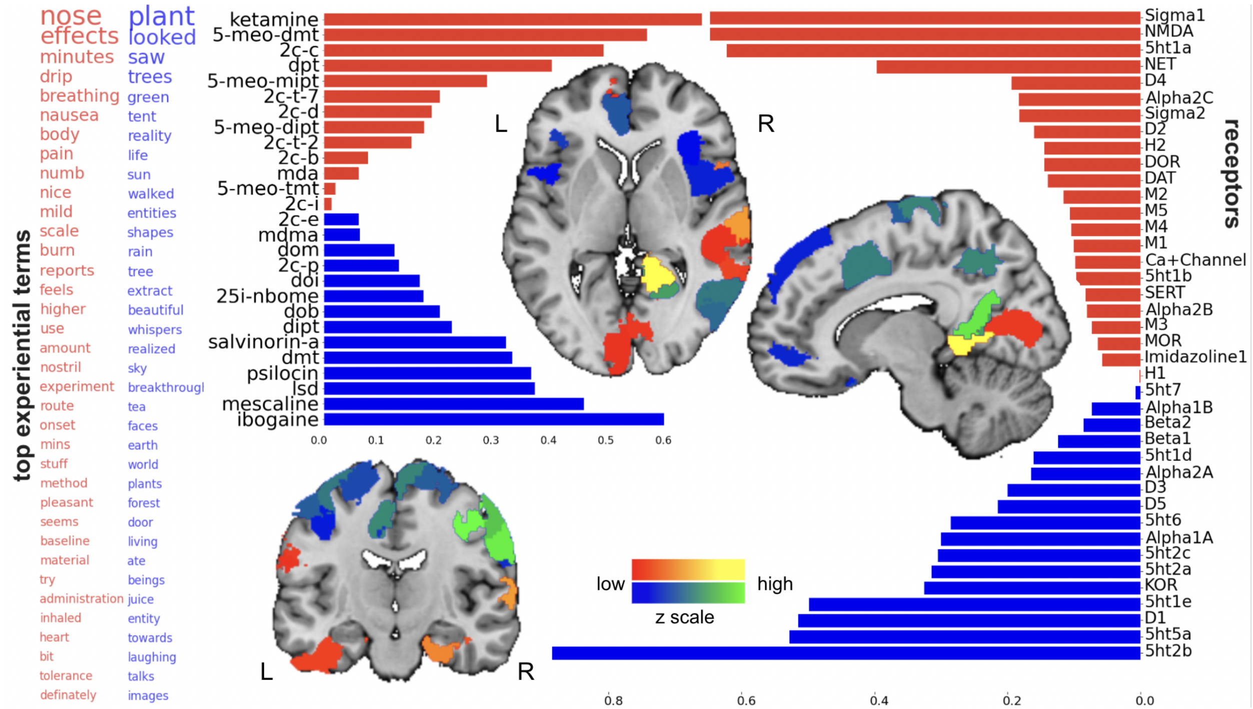


**Supplementary Figure 2: The fifth factor underlying hallucinogenic experiences.** Sagittal, coronal, and axial brain slices are shown at x=11, y=-15, and z=2 (MNI reference space).

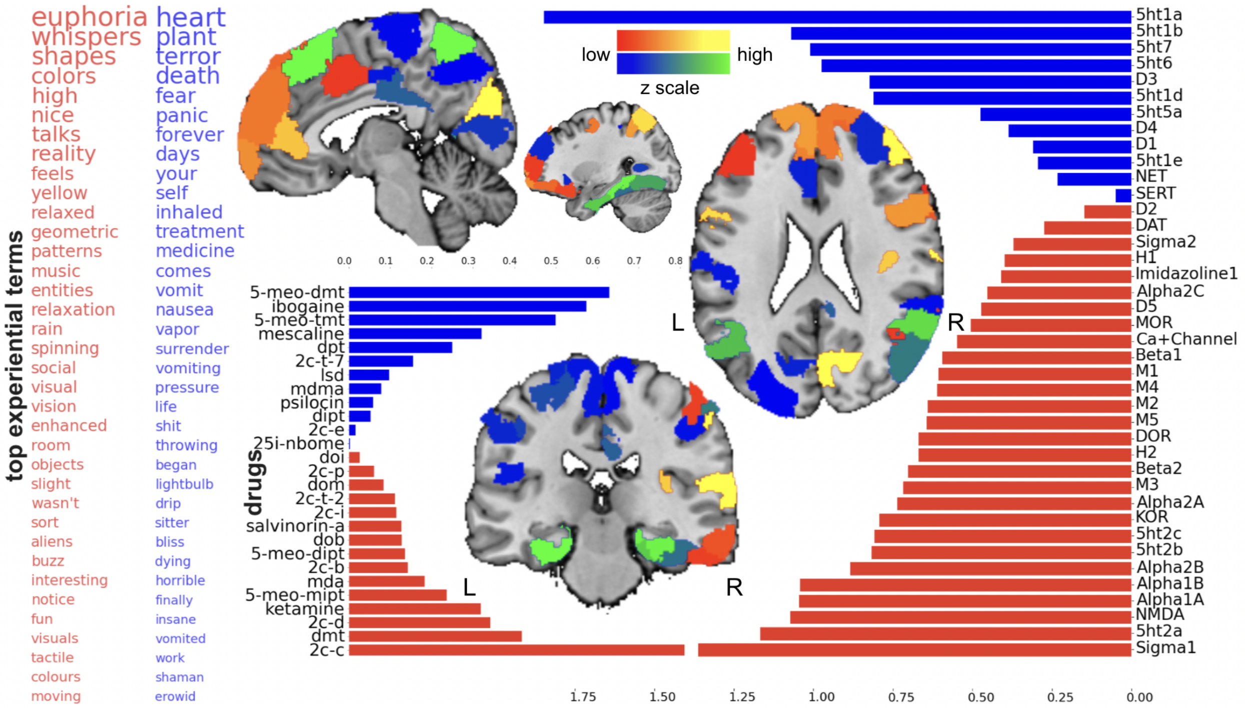


**Supplementary Figure 3: The sixth factor underlying hallucinogenic experiences.** Sagittal, coronal, and axial brain slices are shown at x=5, 28, y=-26, and z=23 (MNI reference space).

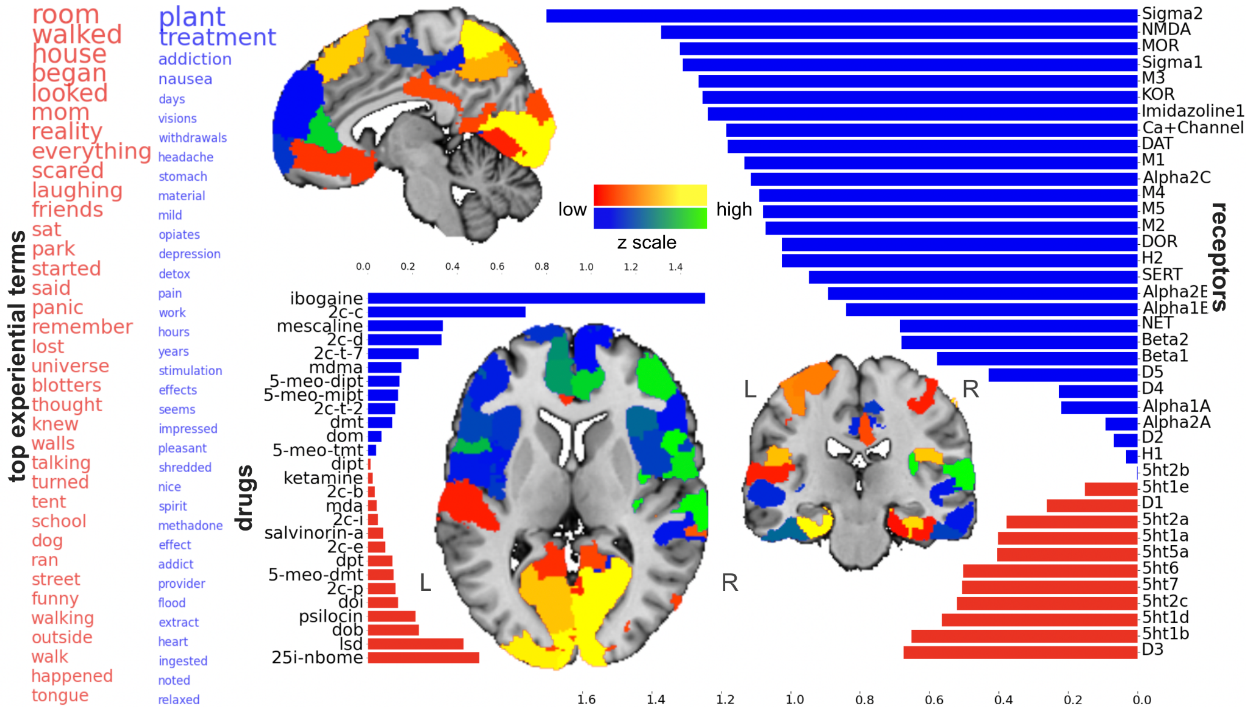


**Supplementary Figure 4: The seventh factor underlying hallucinogenic experiences.** Sagittal, coronal, and axial brain slices are shown at x=7, y=-23, and z=7 (MNI reference space).

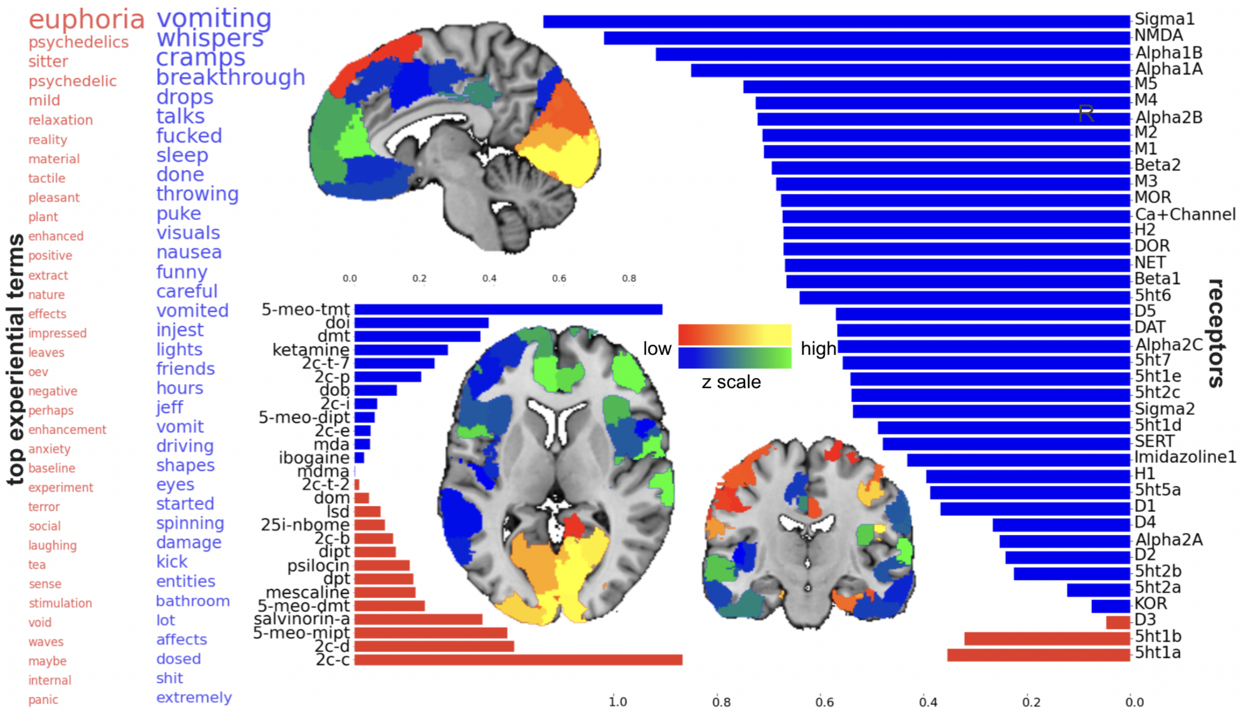


**Supplementary Figure 5: The eighth factor underlying hallucinogenic experiences.** Sagittal, coronal, and axial brain slices are shown at x=-6, y=-17, and z=6 (MNI reference space).
